# MITOCHONDRIAL GENOME REVEALS CONTRASTING PATTERN OF ADAPTIVE SELECTION IN TURTLES AND TORTOISES

**DOI:** 10.1101/2021.02.18.431795

**Authors:** Subhashree Sahoo, Ajit Kumar, Jagdish Rai, Sandeep Kumar Gupta

**Affiliations:** Department of Animal Ecology and Conservation Biology, Wildlife Institute of India, Dehradun; Institute of Forensic Science and Criminology, Panjab University, India

**Keywords:** Testudinoidea, mitochondrial protein-coding genes, OXPHOS subunits, adaptive evolution, natural selection.

## Abstract

Testudinoidea represents an evolutionarily unique taxon comprising both turtles and tortoises. The contrasting habitats that turtles and tortoises inhabit are associated with unique physio-ecological challenges hence enable distinct adaptive evolutionary strategies. To comparatively understand the pattern and strength of Darwinian selection and physicochemical evolution in turtle and tortoise mitogenomes, we employed adaptive divergence and selection analyses. We evaluated changes in structural and biochemical properties, and codon models on the mitochondrial protein-coding genes (PCGs) among three turtles and a tortoise lineage. We used mitochondrial PCGs that constitute the crucial oxidative phosphorylation (OXPHOS) respiratory system, a critical metabolic regulator which assumes key significance in energy regulation of ectotherms.

We detected strong evidence of positive selection along the turtle lineages: Geoemydidae, Emydidae, and Platysternidae, but relatively weak signals in tortoises. The Platysternidae turtles revealed the highest gene and site-wise positive selection. In turtles, positively selected sites were prevalent in NAD2 and NAD4 genes in OXPHOS Complex I, and COB gene of Complex III, indicating convergent adaptive evolution. Besides, NAD3 was the only subunit that showed adaptive selection in both turtles and tortoises, expressing its relevance for all Testudinoidea. Structural and functional analysis revealed many sites and physiochemical changes in important conserved as well as biomedically significant regions, suggesting the influence of adaptive pressure on mitogenome functions. Hence, our study furnished novel evidence of contrasting evolutionary selective pressure acting on closely related groups such as turtles and tortoises with unique habitat preferences and associated eco-physiological challenges.

## 1. Introduction

Global biodiversity continues to be decimated at an unprecedented rate. The challenges threatening sustained survival of the natural species are multi-faceted and often inadequately addressed. Anthropogenic activities have driven a million species towards the risk of extinction with a continually accelerating rate of species decimation **(Tollefson, 2019)**. Despite its severity, anthropogenic pressure that has have driven a million species towards the risk of extinction **(Tollefson 2019)** is a relatively recent phenomenon in the scale of evolutionary development. Natural selection is a long-standing natural pressure that has been insidiously shaping the evolution of species. It regulates species survival and extinction by developing the adaptive potential of species to adapt to a changing environment through micro and macroevolutionary processes **(Harrisson et al., 2014)**. Despite its ubiquity, our understanding of selection pressure on species in need of urgent intervention remains limited. Since selective pressure also acts through microevolutionary tools, its nuances in related species with distinct evolutionary and adaptive needs may often be underestimated. Insight into how natural selection operates on species can be valuable in conceptualizing adaptive conservation strategies **(Eizaguirre and Soares, 2014)**, considering the natural species fitness developed over millions of years.

Chelonians including turtles and tortoises are icons of exemplary evolutionary adaptation. These archaic reptiles have been an important part of the natural ecosystem for about 220 million years and are closely associated with human culture for at least the last 400,000 years **(Stanford et al., 2020)**. Chelonians are perhaps the most easily discernible group with their plastron and carapace, an evolutionary landmark that has compounded their success across an array of ecosystems such as terrestrial, freshwater, and marine. Despite this, more than half of the species of this megadiverse group are threatened with extinction risk due to various anthropogenic pressures **(Stanford et al., 2020)**. Among Chelonians, superfamily Testudinoidea is one of the most endangered and evolutionarily distinct taxonomic units. It comprises four extant families: Geoemydidae (Asian turtles), Testudinidae (Tortoises), Platysternidae (Big-headed turtle), and Emydidae (New World Pond turtles) **(Rhodin et al., 2011)**. Originated in the Cretaceous of Asia, testudinoideans have adapted to a spectrum of habitats and latitudinal range with biological requirements that have led to unique and diversifying physiochemical and morphological specializations **(Lourenco et al., 2012)**.

Often clubbed together from a conservation point of view, turtles and tortoises have shared a 260 million year old evolutionary journey in which tortoises represent a reasonably recent 95 Mya diversification **(Lourenco et al., 2012)**. The morphological and ecological distinctions between turtles and tortoises are a result of divergent metabolic needs arising due to contrasting habitat inhabited by the two groups. These eco-morphological ramifications that make turtles and tortoises fit for a highly aquatic and terrestrial lifestyle, respectively, suggest diverse evolutionary patterns acting on these closely related Chelonians. Evolutionary understanding through natural selective pressure may provide key insight into finer evaluation of strategies governing these unique groups of reptiles.

The counterintuitive forces of dynamic adaptive requirements and functional imperatives act as positive, negative, or neutral selection pressure. Positive selection can be diversifying or directional and enables adaptation, whereas negative selection is purifying and maintaining, which is valuable in eliminating deleterious or harmful substitutions **(Ellegren, 2008)**. Positive natural selection is the key to understand the adaptive potential and hence of scientific interest to uncover the evolutionary process governing species **(Dowling et al., 2008)**. Due to largely uniparental inheritance and lack of recombination, the mitochondrial genome serves as an apt model to study patterns of natural selection. The most vital function of the organelle is to generate cellular energy through the critical OXPHOS system **(Das, 2006)**.

The respiratory complexes in the OXPHOS respiratory system are under the dual control of both mitochondrial and nuclear genomes **(Bayona-Bafaluy et al., 2005)**. The 13 PCGs in the mitochondrial genome are components of four of the five respiratory complexes in the OXPHOS biochemical pathway: complex CI (seven NADH dehydrogenases: NAD1-NAD6, NAD4L), CIII (Cytochrome *b*; COB), CIV (three cytochrome c oxidases subunits: COI, COII, and COIII) and CV (two ATPase: ATP6 and ATP8). Through the OXPHOS biochemical pathway, electrons are transferred from one complex to another to be reduced at CIV **(Gu et al., 2019)**. As electrons proceed further down the system, protons are pumped into the intermembranous space creating an electrochemical gradient. ATP synthase in CV uses this gradient energy to synthesize ATP **(Ludwig et al., 2001)**. Cellular respiration is the primary source of energy and significantly influences metabolic performance, which has implications for species with unique and diverse metabolic needs **(Shen et al., 2009)**.

Despite natural selection on mitochondrial PCGs being primarily purifying to preserve crucial gene functions, adaptive novelty may induce advantageous amino acids (AAs) substitutions **(Dowling et al., 2008)**. Increasing evidence of positive selection on mitochondrial OXPHOS complex genes linked to physiochemical adaptations in response to metabolic needs has been previously reported in studies, such as larger brains in anthropoid primates **(Doan et al., 2016)**, bats and flight of origin **(Shen et al., 2010)**, killer whales in the Antarctic **(Foote et al., 2011)**, polar bear and adaptation to Arctic environment **(Welch et al., 2014)**, swimming performance in fishes **(Zhang and Broughton, 2015)**, and scute loss in turtles **(Escalona et al., 2017)**. This suggests that diverse taxa cope with numerous metabolic challenges demanding ingenious morphological and physiochemical adaptations in response to environmental needs. Variations in mitochondrial PCGs have been associated with novel thermo-metabolic demands in various species under environmental adaptive opportunities **(Da Fonseca et al., 2008; Garvin et al., 2015)**. With these novel evidences, we hypothesized that natural selection might be acting differently on tortoises and turtles.

To test our hypothesis of molecular adaptation acting differently on turtles and tortoises, we measured the pattern and intensity of natural selection acting on mitochondrial PCGs. We tested lineage-specific positive selection employing two widely used methods: changes in physicochemical properties using a phylogenetic tree **(McClellan and McCracken, 2001)** and maximum likelihood (ML) codon-based models estimating non-synonymous to synonymous substitution rate ratio (ω) **(Goldman and Yang, 2004)**. The ω value suggests the mode and intensity of selection, where ω < 1 indicates negative selection; ω = 1 represent neutral selection, and ω > 1 shows positive selection. We also inferred the functional relevance of positively selected sites (PSS) in OXPHOS protein complexes through protein modeling. Based on our findings, we discuss the comparative adaptive evolution in turtles and tortoises associated with metabolic demands in biochemistry and physiology.

## 2. Results

### 2.1. Phylogeny estimates

We constructed the phylogenetic relationship among Testudinoidea employing Bayesian and Maximum Likelihood (ML) methods using concatenated sequences of the 12 PCGs **(Fig. 1)**. The sequence of Leatherback sea turtle (*Dermochelys coriacea*: accession number MF460363) was chosen as the outgroup. Both methods yielded similar tree topologies. Bayesian-based high posterior probability (PP >70) and ML bootstrap proportions (BP>0.60) were represented at respective nodes. Both the phylogenetic trees were congruent with accepted turtle phylogenies based on extensive mitochondrial and nuclear DNA, as well as fossil evidence **(Lourenco et al., 2012; Goldman and Yang, 1994)**.

**Figure 1.**
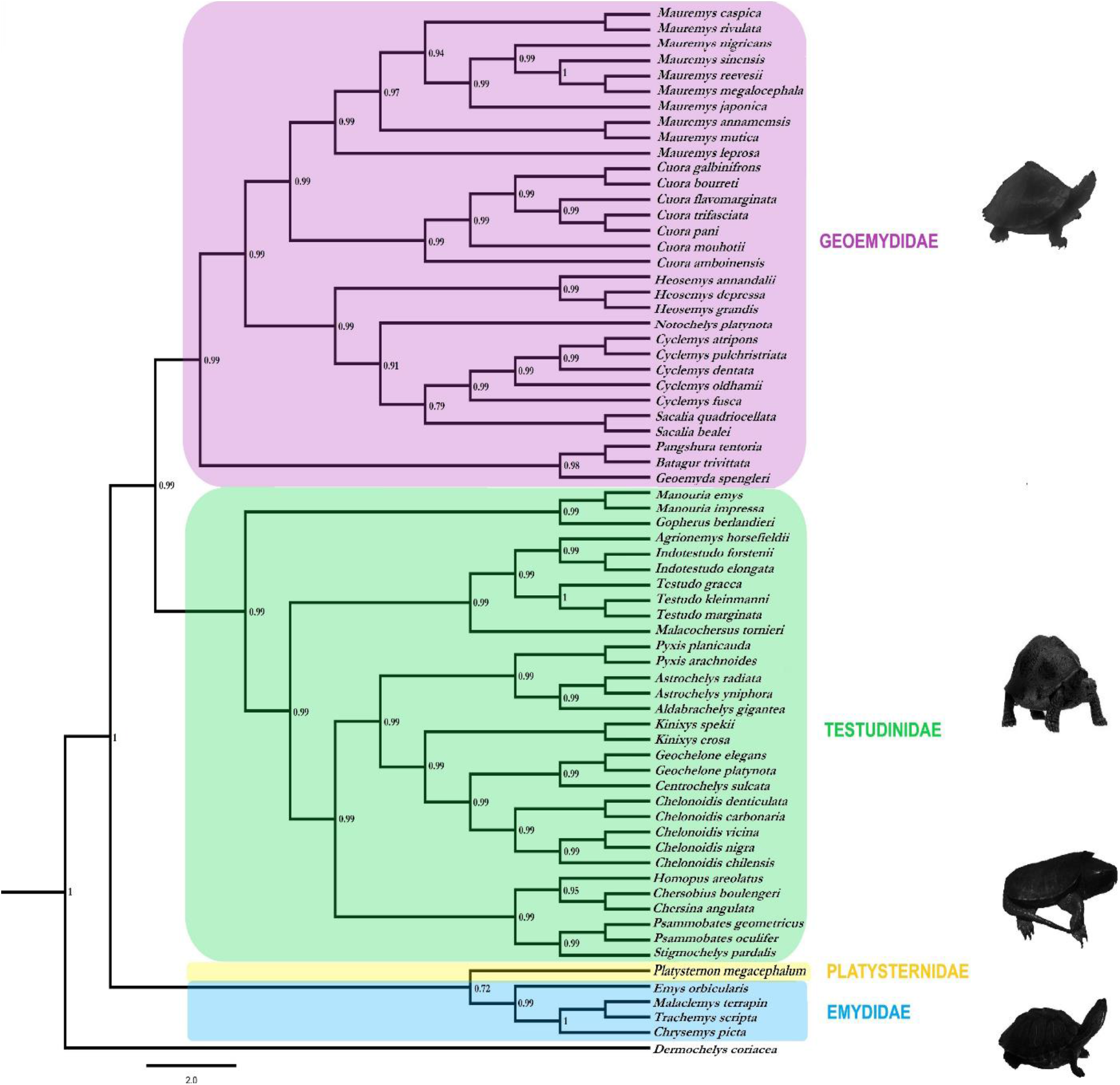
Bayesian phylogenetic tree for Testudinoidean turtles and tortoises. The tree was constructed with 12 PCGs’ concatenated sequences (with *Dermochelys coriacea* as outgroup). Branch lengths and scale bar indicate nucleotide substitutions per site. Numbers next to nodes represent bootstrap probability values for 1000 runs.

### 2.2 Adaptive selection and divergence in tortoise versus turtles

The TreeSAAP and CODEML (BrS) analysis revealed contrasting rate of adaptive selection acting on turtles as compared to tortoises. In Testudinidae, at the NAD2 gene of CI, a single site 85 with AAs substitution S→D was detected under positive selection, which led to an increase in Polar requirement (Pr) and αc **(Fig. 2)**. In contrast, overall, nine genes with 41 sites were found to be under positive selection in turtle families: Geoemydidae, Platysternidae, and Emydidae **(Table 1; Supplementary Table: ST2)**. Several significant AAs substitutions associated with changes in physicochemical properties under adaptive selection were detected in turtles across CI, III, IV, and V **(Supplementary Table: ST3, ST4)**.

**Figure 2.**
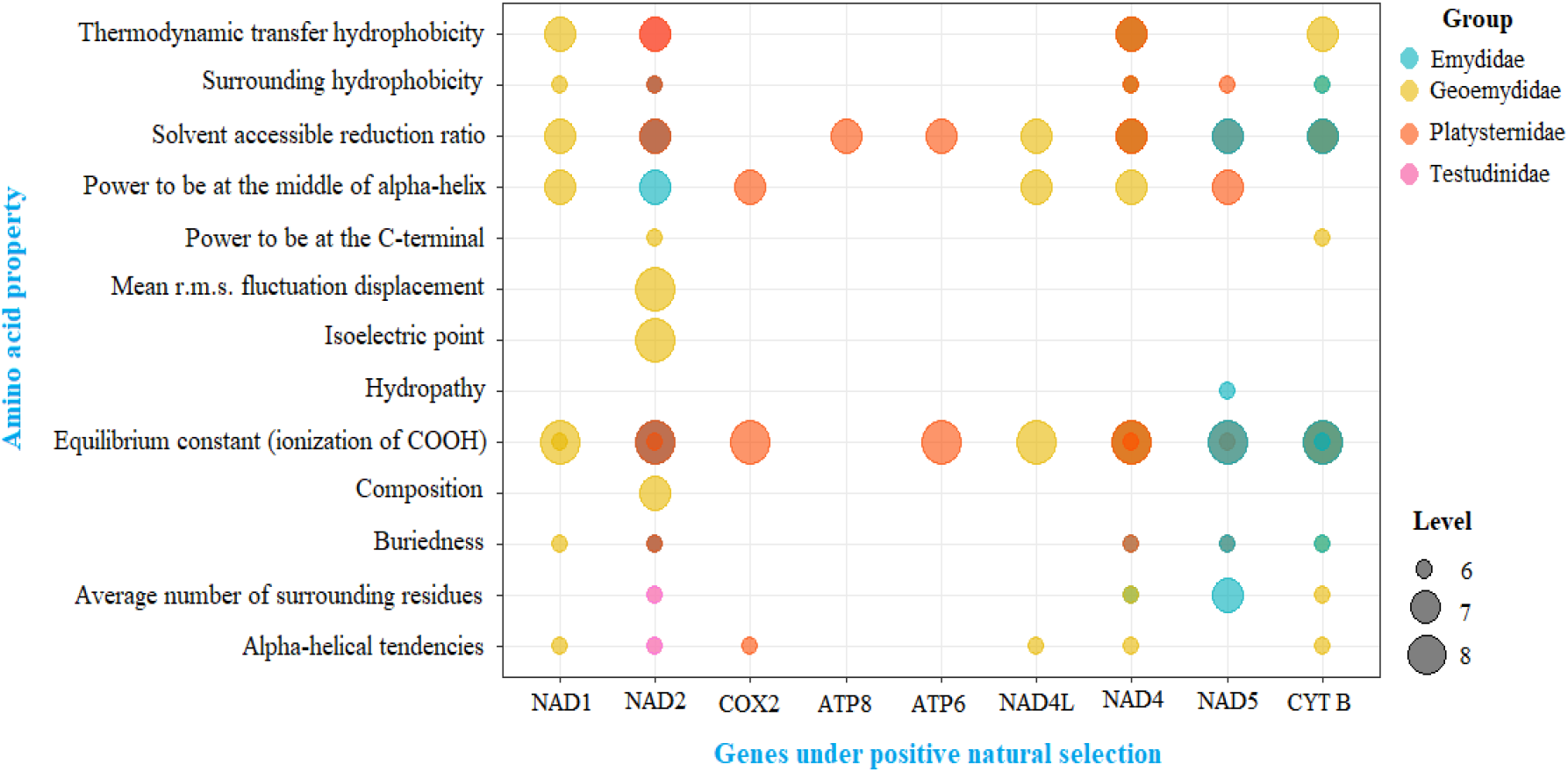
Distribution of change in AAs properties with magnitude 6 to 8 associated with AAs substitutions at PSS across genes in specific turtle and tortoise families.

**Table 1.**
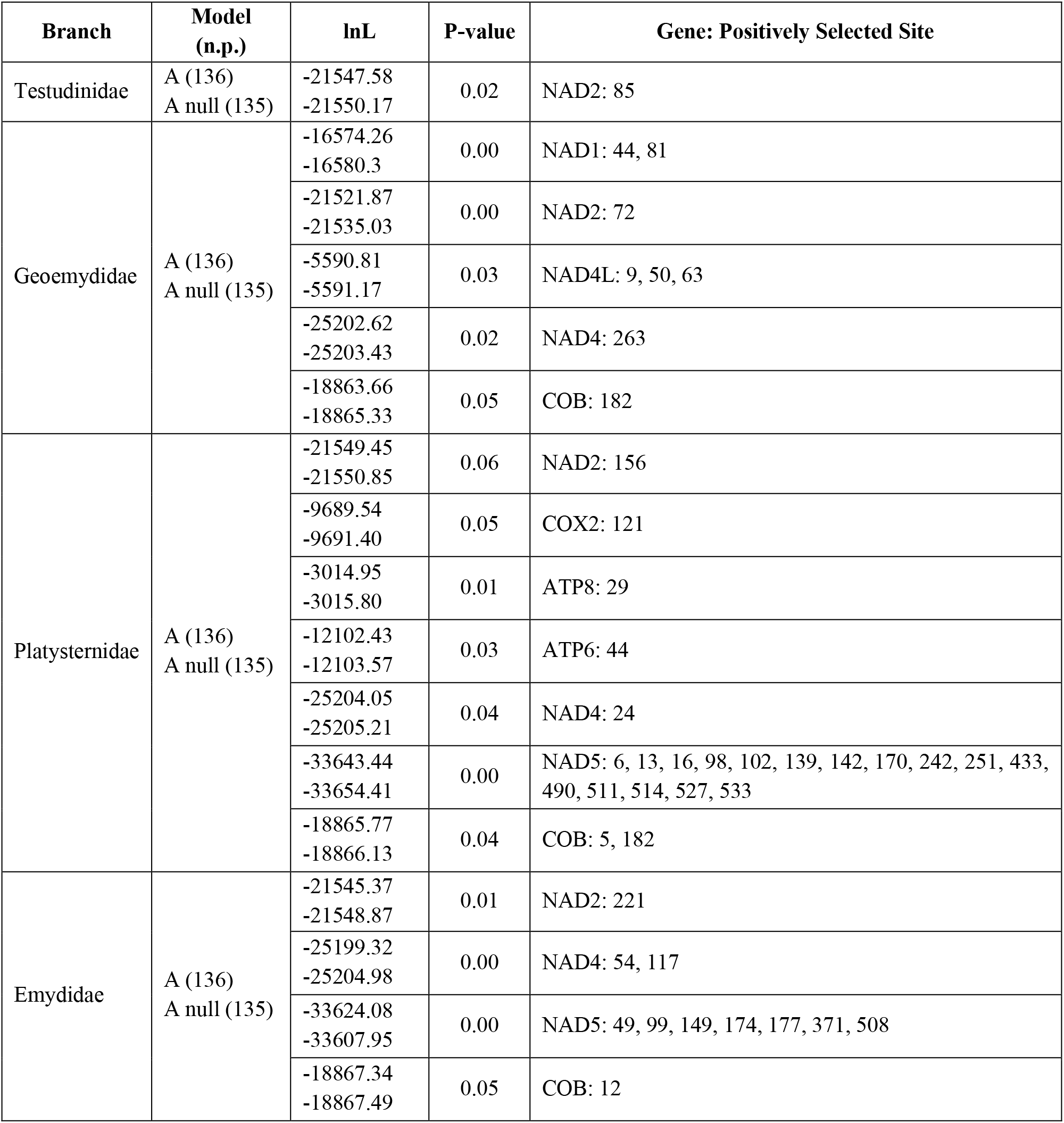
Branch-site (BrS) model analyses of the gene-specific mtDNA data sets: Sites identified as positively selected with Bayes Empirical Bayes (BEB) method having Posterior Probability (PP) >= 0.90 are listed. Complete results including ω estimates for site classes with support values are provided in the Supplementary file: ST3; n.p.: number of parameters.

In Geoemydidae, a total of eight PSS were detected out of which seven sites (2 in NAD1, 1 in NAD2, 1 in NAD4, and 3 in NAD4L) were located in CI, and only one site was found in COB gene of CIII. We detected changes in 17 physicochemical properties associated with those PSS. In Platysternidae, 18 sites were found in CI (1 in NAD2, 16 in NAD5 and 1 in NAD4), two sites in COB of CIII, one site in COX2 of CIV and two sites in CV (1 each in ATP6 and ATP8) under positive selection. Interestingly, the site 182 in the COB gene was found under positive selection in both Geoemydidae and Platysternidae. We detected changes in 12 physicochemical properties in Platysternidae, indicated to be under significant fluctuation in the PSS. In Emydidae, ten sites (1 in NAD2, 7 in NAD5 and 2 in NAD4) of CI and a single site in COB of CIII were found to be under positive selection with changes across seven physicochemical properties.

For selection analysis, LRTs were significant (P < 0.05) for all genes under positive selection. Among them, PSS in the NAD2 gene was found to be common across both turtles and tortoises, whereas the sites found in NAD4 and COB genes were only detected in turtle species. Significant changes in physicochemical properties and their higher occurrence in turtles as compared to tortoises signifies accelerated positive selection associated with novel substitutions in turtles. Based on our comparative analysis of turtle and tortoise, we found natural selection acting differently on turtles and tortoises with evidence of higher adaptive selection in turtles as compared to tortoises.

### 2.3. Adaptive selection and divergence across Testudinoidea

Out of 31 physicochemical properties analyzed by TreeSAAP in the NAD3, overall, 17 were detected in Testudinoidea, which includes 7 in Emydidae, 11 in Platysternidae, 14 in Geoemydidae and 14 in Testudinidae **(Supplementary Table: ST5)**. 17 unique AAs substitutions types were represented by 62 substitutions, which were significantly (P<0.005) detected under adaptive pressure. The results revealed that Testudinidae contained 30 AAs substitutions, which was highest among the Testudinoidea. Testudinidae shared 12 AAs substitutions with Geoemydidae, followed by five and three AAs substitutions with Emydidae and Platysternidae, respectively. It was congruent with the phylogenetic tree topology, where the Geoemydidae was the sister group and closely related to Testudinidae.

The random-sites codon model detected pervasive patterns of adaptive evolution in Testudinoidea PCGs. ω and kappa (κ) estimates in the M0 model implemented in CODEML varied across gene-complexes confirming non-uniform rates of evolutionary selection acting on mitochondrial PCGs in the OXPHOS system **(Table 2)**.

**Table 2.** Gene wise-M0 codon model: The M0 codon model was implemented for each of the 12 PCGs’ alignment to estimate omega (ω), kappa or the transition to transversion ratio (κ); the sequence length in codons is provided along with the gene wise log-likelihood values (lnL) and the number of non-synonymous substitutions per site (dN).

| Genes | Sequence length | lnL | Branch specific dN Estimates | | $\omega$ | $\kappa$ |
| --- | --- | --- | --- | --- | --- | --- |
|  |  |  | Range | Median |  |  |
| **NAD1** | 310 | -16765.27 | 0-0.0496 | 0.0248 | 0.06 | 12.428 |
| **NAD2** | 346 | -21670.97 | 0-0.0773 | 0.03865 | 0.05 | 24.610 |
| **COX1** | 492 | -21806.19 | 0-0.0351 | 0.01755 | 0.03 | 15.060 |
| **COX2** | 220 | -9814.09 | 0-0.0325 | 0.01625 | 0.05 | 16.100 |
| **ATP8** | 48 | -3083.24 | 0-0.2368 | 0.1184 | 0.26 | 12.923 |
| **ATP6** | 220 | -12279.72 | 0-0.0811 | 0.04055 | 0.10 | 13.916 |
| **COX3** | 245 | -10768.64 | 0-0.0373 | 0.01865 | 0.04 | 16.483 |
| **NAD3** | 104 | -6844.41 | 0-0.1286 | 0.06515 | 0.66 | 10.245 |
| **NAD4L** | 95 | -5670.59 | 0-0.0679 | 0.03395 | 0.11 | 10.991 |
| **NAD4** | 440 | -25635.02 | 0-0.0766 | 0.0383 | 0.07 | 16.040 |
| **NAD5** | 560 | -33997.10 | 0-0.1739 | 0.08695 | 0.11 | 12.050 |
| **COB** | 366 | -19117.58 | 0-0.0583 | 0.02915 | 0.05 | 13.432 |

Among gene complexes, CIII and CIV, comprising COB and CO*I*, CO*II* and CO*III*, respectively, were under the strongest purifying constraint with lowest ω estimates, ranging from 0.03-0.05. In contrast, the gene NAD3 of CI and ATP6 and ATP8 of CV were found to be under lower purifying constraints with higher ω estimates, ranging from 0.10-0.66. Considerable variation among various gene complexes indicated that selective pressure was not equal on all genes or associated OXPHOS complexes. Furthermore, to detect sites under positive selection across Testudinoidea, M8a-M8 nested models were implemented with Likelihood Ratio Tests (LRTs). The site-specific posterior estimates of dN/dS estimates showed variation and intensity of selective pressure acting on various PCGs **(Fig. 3)**.

**Figure 3.**
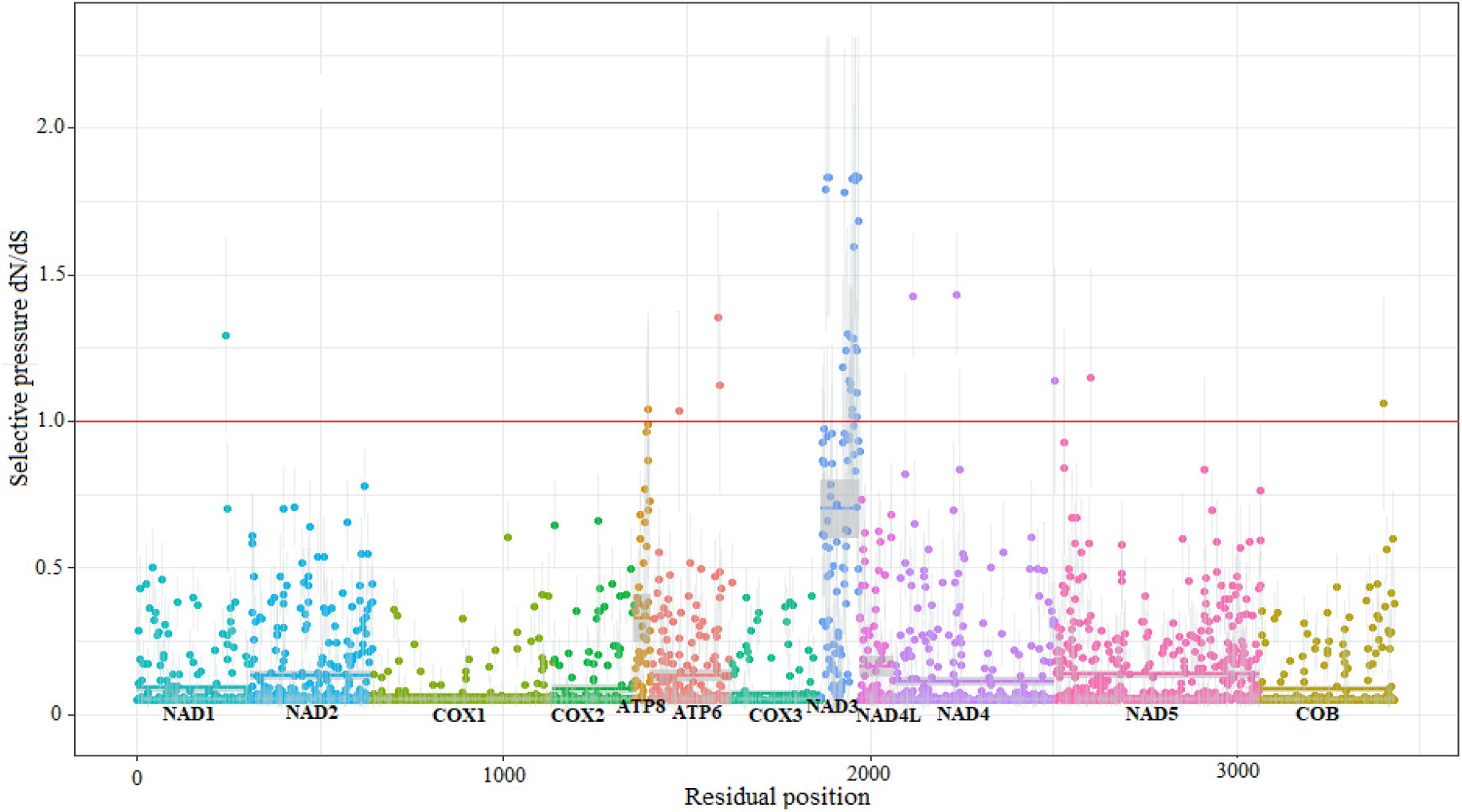
Gene wise estimates of dN/dS for the concatenated data set of 12 mitochondrial PCGs were calculated using M8a-M8 nested random-site models. The y-axis denotes the BEB posterior mean estimate of dN/dS for each site with grey vertical bars indicating standard error. Horizontal solid lines indicate mean estimate of the genes.

The order of the estimates of dN/dS were NAD3 > NAD4 > ATP6 > NAD1 > NAD5 > COB > ATP8. The nested model test also detected significant PSS in the NAD3 region of CI. A total of eight sites (13, 16, 19, 63, 84, 92, 94, and 101) were detected with BEB (PP>95%) and significant LRTs (P<0.001) **(Supplementary Table: ST6)**. This pattern was congruent with selective pressure detected under the M0 model. Thus, the NAD3 gene in CI was found to be the only region under adaptive selection in tortoise (Testudinidae) and turtle (Geoemydidae, Platysternidae, and Emydidae) indicating its important role in shaping the adaptive pattern for all Testudinoidea.

### 2.4. Structural and functional divergence analysis

Both TreeSAAP and CODEML identified that the NAD3 region was under adaptive selection in tortoise and turtles, containing eight PSS with significant changes in AAs: 13I, 16T, 19I, 63P, 84Y, 92T, 94N, and 101S. Geoemydidae showed similar sites with changes in AAs under adaptive selection, as found in Testudinidae. In Emydidae, AAs substitutions were present at six sites: 13, 16, 19, 92, 94, and 101, whereas in Platysternidae, five AAs substitutions were present at sites: 13, 84, 92, 94 and 101. The changes in physicochemical properties were significantly selected for magnitude >6.

Specific AAs substitutions detected by TreeSAAP in the NAD3 region were located on three different Transmembrane Helix (TMH) viz, sites 13, 16, and 19 were found on TMH1, 63 on TMH2 and 84, 92, 94 and 101 on TMH3 **(Supplementary Table: ST8)**.

Site 13 was associated with T→I substitution in Testudinidae, Emydidae, and Platysternidae with an increase in solvent-accessible reduction ratio (Ra) and Thermodynamic transfer hydrophobicity (Ht) across all branches. Platysternidae had an additional increase in Equilibrium constant of ionization (pK) associated with the same substitution. In contrast, the substitution of T→A and T→V was only found in Testudinidae with an increase in α-helical tendencies (Pα) and Ra, respectively. The AAs substitution of A→V at site 13 was only detected in Geoemydidae with an increase in β-structure tendencies (Pβ). Site 16 showed AAs substitution shared by both Testudinidae and Geoemydidae. While the change from T→A led to an increase in Pα in both branches, the substitution from T→I was associated with an increase in Ra, Ht and pK only in Geoemydidae. At position 19, the substitution of T→I was observed in Testudinidae and Geoemydidae, which was similar to substitution at sites 13 and 16 with an increase in Ra and Ht.

Additionally, reverse AAs substitution was also observed I→T in Testudinidae and Geoemydidae, which led to a decrease in Ra and Ht. A significant increase in pK was also seen with two other AAs substitutions in Testudinidae, T→V and I→M with the latter being shared with Emydidae. Interestingly, in TMH2 AAs, substitution I→V at site 63 led to a drastic increase in pK, which was unique in Testudinidae. At site 84 on TMH3, AAs substitution S→P was observed in both Testudinidae and Geoemydidae with an increase in Coil tendencies (Pc) and power to be at C-terminal of α-helix (αc), respectively. The reverse AAs substitution with a decrease in Pc was observed only in Testudinidae.

Site 92 showed L→P substitution in Testudinidae, Geoemydidae, and Platysternidae, which led to a shared steep increase in Pc and αc, whereas a decrease in αn and αm was only observed in Testudinidae and Geoemydidae. In contrast, Platysternidae showed an increase in power to be at the N-terminal of α-helix (αn). In Testudinidae, an additional drastic increase in Pc and αc was observed. In the same position, P→S was detected both in Testudinidae and Emydidae with a decrease in αc and αm only observed in Testudinidae. Additionally, Testudinidae underwent a unique AAs substitution at this site of P→L with a decrease in compressibility (K0), Mean r.m.s. Fluctuation displacement (F) and Turn tendencies (P) accompanied by a sharp decrease in Pc, αc, and power to be at the middle of α-helix (αm). At Site 94, AAs substitution Y→H was observed only in Geoemydidae that led to an increase in αm. The AAs substitution P→A at site 101 led to an increase in Pα and αn with a decrease in P and Pc, and S→F led to an increase in Ht with a decrease in P and F, in Testudinidae. P→L was another unique substitution detected in Testudinidae at this site with a drastic increase in αn and a decrease in P, F, and K0 and a steep decrease in αc and Pc. The AA substitution of P→F was detected in both Testudinidae and Geoemydidae with identical changes in physicochemical properties: increase in Ra but decrease in P, F, K0, and Pc.

### 2.5. Structural analyses

PSS identified using TreeSAAP and CODEML through both random and branch-site analyses were projected onto crystallized structures of mitochondrial PCGs in the context of contemporary structural and biomedical insights. Considering only sites with PP > 0.90, a total of 43 PSS was mapped onto 3D protein structures. Many of these sites were found to be located close to the key residue and conserved regions, suggesting their substitutions to be adaptively significant **(Fig. 4)**.

**Figure 4.**
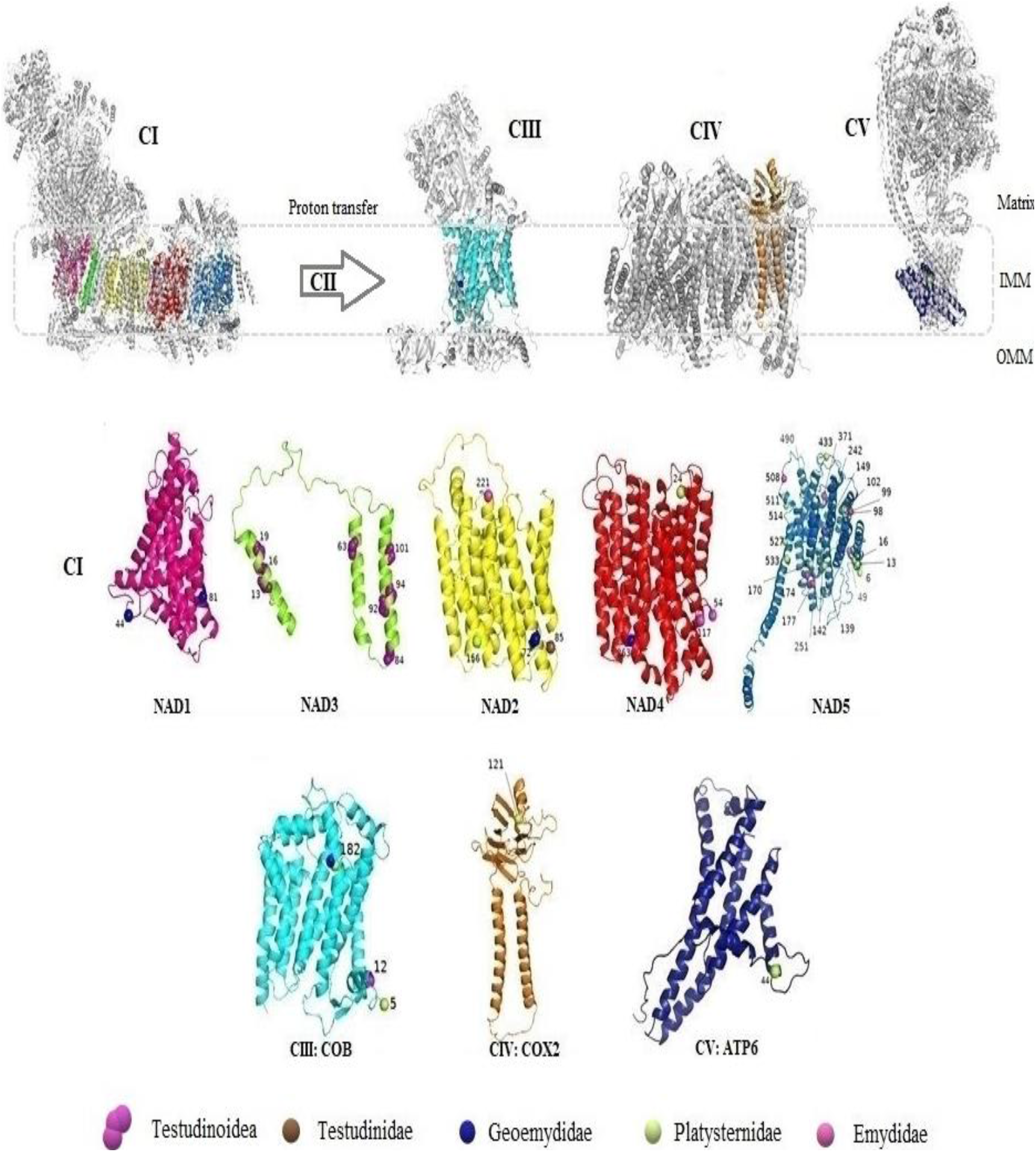
Graphical representation denoting the distribution of PSS in Testudinoidean lineages across the OXPHOS system. All the 12 mitochondrial PCGs encoding the OXPHOS system were projected onto homologous protein structures and are represented in different colours. Grey structures represent the entire OXPHOS complex and no solved crystallized structures were available for the ATP8 gene. The NAD3 gene (lemon green) represents sites under positive selection across all Testudinoidean lineages as detected by random sites analysis. The rest of the genes, NAD1 (hot pink), NAD2 (yellow), NAD4 (red), NAD5 (cobalt blue), COB (cyan), COX2 (orange) and ATP6 (deep blue) indicate sites under lineage-specific positive selection detected by Branch site analysis. The four Testudinoidean lineages are represented by singular spheres of different colours: Testudinidae (brown), Geoemydidae (dark blue), Platysternidae (lemon yellow) and Emydidae (pink). The Testudinoidean positive sites are indicated by multiple pink spheres in NAD3. Only PSS with PP > 0.90 and significant LRTs were considered. IMM: Inner mitochondrial membrane; OMM: Outer mitochondrial membrane.

## 3. Discussion

Our analysis revealed high adaptive selection rates in turtle mitogenome compared to low adaptive pressure on tortoise mitogenome that borderline purifying. There may be a few possible explanations for such contrasting adaptive pressure. Firstly, adaptive selection may indeed be influencing the tortoise genome, but mostly through nuclear subunits, which were not accommodated in our study. Secondly, tortoises due to their unique physiochemical and metabolic requirements, may have been under long-standing stabilizing divergent selections similar to patterns of adaptation observed in fishes and cetaceans **(Boeuf and Payan, 2001; Wang et al., 2015)**. Lower metabolic requirements, higher gestation period, characteristically high longevity, lower speciation rate and adaptation for an arid terrestrial life might have brought about enough morphological, ecological, and metabolic adaptations, which were sufficient for their habitats and energetic demands without significant adaptive adjustments in the OXPHOS subunits **(Avise et al., 1992; Rodrigues and Diniz-Filho, 2016)**. Lastly, positive selection may have been rapid and episodic in tortoises, followed by a relatively long purifying selection, which might have masked signatures of positive selection.

We performed TreeSAAP and BrS analysis to examine the patterns of positive selection in turtles and compared it to tortoises, additionally random site analysis was also incorporated to understand selective pressure on all Testudinoideans. The results of BrS revealed numerous sites under positive selection in turtles compared to weaker positive selection in tortoises. In contrast, the random site analysis revealed predominantly purifying selection acting on the mitogenome of Testudinoidea with a single gene (NAD3) identified under positive selection. It highlighted the adaptive significance of the NAD3 gene for all Testudinoideans. The NAD3 subunit is relatively small and one of the core subunits in CI, which, together with nuclear-encoded subunits, makes the proximal pump module that links the matrix arm with the membrane arm OXPHOS system **(Wu et al., 2016)**. It also holds the quinone binding site at its interface with NAD1 and is less conserved compared to other subunits, enabling more adaptive flexibility **(Fiedorczuk et al., 2016)**. It is noteworthy that most sites associated with disease causing substitutions have been identified in the loop of NAD3 **(Bridges et al., 2011)**, while all the sites detected to be under positive selection in our analysis were located in one of the three TMH.

### 3.1. Mitogenome adaptive evolution in turtle versus tortoise and its biochemistry

In all Testudinoidea, the NAD3 was the most prominent gene detected under positive selection and hence attested to adaptive divergence among the branches. Interestingly, a drastic decrease in PSS's significant physicochemical properties was observed in Testudinidae across sites 19, 84, 92 102, present in TMH1 and TMH3. Preference for N-cap in the α-helix interior over C-terminal and middle was observed in Testudinidae **(Cochran and Doig, 2001)**. Reduction in the solvent-accessible reduction ratio may cause decrease in space for active site formation **(Ponnuswamy and Gromiha, 1993)**. Decrease in important structural properties such as the side chain's hydrophobic character, coil tendencies, compressibility, and turn tendencies were observed **(Marcelino and Gierasch, 2008)**. These results indicate the crucial role of these AAs properties in the divergent evolution of Testudinidae through structural and biochemical changes. Thus, the analysis revealed large scale variation in adaptive pressure and substitution pattern across all branches, indicating varying natural selection between turtles and tortoises. Hence, we further explored branch-specific selective pressure comparing both turtles and tortoises.

The result of our analysis suggested that adaptive selection has acted upon multiple genes across OXPHOS complexes in turtles but has had largely purifying constraint on tortoise mitogenome. Here we discuss focusing on a few sites with physiochemical and structural significance. Among the OXPHOS, CI was the only complex that contained sites under positive selection in both turtles and tortoises. While turtles’ mitogenome revealed positive selection across a total of 36 sites across five genes: NAD1, NAD2, NAD4, NAD4L, and NAD5 in CI, the tortoises’ mitogenome had a single positively selected site in NAD2. CI or the NADH: Ubiquinone oxidoreductase powers ATP production in mitochondria by reducing NADH potential moving regulating channeling protons across the inner mitochondrial membrane **(Zhu et al., 2016)**. CI was found to be the gene-complex with the maximum number of sites and genes under positive selection. NAD2 and NAD4 were the only genes in CI to contain PSS in all turtle branches. NAD4 is a part of the distal antiporter like proton pump module at the end of the membrane arm of CI, causing conformational change in the complex that accommodates binding of CIII and CIV in the respiratory pathway **(Gu et al., 2019)**. These findings were congruent with meta-analysis of numerous species **(Garvin et al., 2015)** and have been previously reported in fishes **(Consuegra et al., 2015)**, turtles **(Escalona et al., 2017)**, and mammoths **(Ngatia et al., 2019)**, that have highlighted CI as an important tool for rampant adaptive selection.

In turtles, Geoemydidae was the only family where PSS was detected in NAD1 and NAD4L genes. The site 44 in the NAD1 gene was found to be a functionally conserved residue located in the Inter Transmembrane Helix (ITMH) between TMH1 and TMH2. We observed changes in physicochemical properties associated with hydrophobicity and folding rate that may have significant implications on protein structure and function **(Mallik et al., 2016)**. In the NAD4L gene in Geoemydidae, three sites were detected under positive selection. Noteworthy, site 63 was in a pathologically relevant region associated with LHON disease **(Mitchell et al., 2006)**.

In NAD2 and NAD4 of CI, unique PSS sites were detected in each family of turtles. In the NAD2 region, site 221 of Emydidae, located in the TMH8, was found to be highly conserved and involved in ion transport and catalysis **(Sazanov, 2015)**, whereas site 156 in Platysternidae was situated in a highly conserved region in THM6. Interestingly, site 72 in Geoemydidae was located in the TMH3 region associated with the substitution causing Leigh disease **(Ugalde et al., 2007)**. In addition to this, the single-site 85 in NAD2 is in the ITMH coil between TMH3 and TMH4, both of which are critical conserved regions with essential sites of proton translocation pathway **(Fiedorczuk et al., 2016)**. Surprisingly, in the NAD2 region, positive selection sites were detected in all branches of turtles and tortoises, but it was not detected under Random site analysis. It suggested that the NAD2 gene had adaptively evolved in a branch-specific manner across all Testudinoidea. In contrast, NAD3 was detected to be under positive selection in Random site analysis and not in BrS analysis. This suggested that NAD3 sites might have undergone adaptive selection in a widespread manner and not branch-specific, unlike NAD2. Moreover, in the NAD4 gene, PSS was detected only in turtles, and all the sites were located on THM conserved regions. The site 117 detected in Emydidae was located in a highly conserved region of TMH5 **(Bridges et al., 2011)**.

The NAD5 gene comprised the greatest number of PSS but was detected only in Emydidae and Platysternidae. Of the seven PSS in Emydidae, site 177 in TMH6 was detected to be a conserved residue with substitutions leading to potentially significant physiochemical consequences. Out of the sixteen PSS detected in Platysternidae, sites 102 and 142 were located in highly conserved regions of TMH3 and THM5, respectively. Additionally, site 251 was located near residues where AAs substitutions have been found to be associated with CI related disorders such as Ataxia in Humans **(Valente et al., 2008)**.

The CI is the point of entry for electrons into the OXPHOS system, while CIII and CIV work in sync with CI to coordinate electron transfer with proton translocation across the inner mitochondrial membrane **(Gu et al., 2019)**. Like NAD2 and NAD4 genes, the COB gene of CIII or Cytochrome bc1 complex was found to be under positive selection in all turtles. The COB gene is a highly conserved region of the OXPHOS system with a pivotal role in mitochondrial energy generation. It catalyzes the reversible transfer of electrons from ubiquinol cytochrome c to be used in proton translocation **(Howell, 1989)**. In contrast to tortoises, where the highly conserved CIII is maintained under purifying selection owing to its vital metabolic role, its adaptive evolution in turtles signifies the genes’ adaptive potential to cope with energetic demands unique to turtles. CIV is at the distal end of the OXPHOS complex, catalyzing the oxidation of reduced cytochrome c and oxygen molecules to water **(Belevich et al., 2007)**. ATP synthase in CV uses the proton gradient from H+ pumped into the intermembranous space to generate ATP, which requires significant conformational changes in the membrane arm **(Fillingame at al., 2003)**. CIV and CV were found to be under adaptive pressure only in Platysternidae, highlighting unique evolutionary adaptations in the species.

Most of the sites under positive selection were found to be located in the TMH of specific proteins. Radical changes in AAs properties mostly led to substitutions involving residues associated with the interior of folded protein structure as compared to those exposed to solvents. It suggested exposed residues to be under greater selective constraint as compared to interior residues. AAs substitution coupled with BrS analysis, revealed convergent evolution of turtle mitogenome through NAD2 and NAD4 genes in CI and COB gene in CIII. Adaptive evolution in these structural gene complexes may have influenced the efficiency of respective respiratory chain complexes shaping higher adaptive potential in turtle mitochondrial OXPHOS system compared to tortoises.

### 3.2. Physio-ecology comparison of turtle versus tortoise

Turtles and tortoises have evolved as cold-blooded reptiles with distinctly efficient aerobic capacity. In the evolution of ectothermy, mitochondrial ATP generation and metabolic efficiency are both strongly influenced by temperature. It has been found that mitochondrial thermal physiology may vary across ecology and thermal flexibility **(Clarke and Portner, 2010)**. Tortoises are slow reptiles that thrive in desert, arid grasslands, scrub to wet evergreen forests, and have distinguishable clumped toes fit for terrestrial lifestyle **(Vitt and Caldwell, 2013)**. In contrast, an aquatic lifestyle in turtles comes with additional energy demands that require a highly efficient metabolic system for which all Testudinoidea turtles are adept with webbed toes. Compared to the widely distributed tortoises, most turtle families are limited to specific geographic habitats, which could have encouraged distinct adaptive adjustments. In terms of diet, tortoises are restricted mostly to the herbivorous end of the spectrum, while turtles span across from strictly herbivores to strictly carnivores, expanding their nutritional options **(Rodrigues and Diniz-Filho, 2016)**. Moreover, added energy investment into heavier shell morphology of tortoises is negatively associated with energy metabolism as well as its activity levels **(Zhang et al., 2019)**.

Higher adaptive potential in aquatic turtles mitogenome compared to terrestrial tortoises could also be associated with contrasting habitats and diversification patterns. Habitat acts as a vital determinant of ecological and physiological adaptations. Along with ecological opportunities, island invasions, and climatic factors, habitat drives diversification and species evolution **(Rodrigues and Diniz-Filho, 2016)**. Unlike several vertebrates where higher speciation rates were observed in terrestrial species, turtles exhibit a higher rate of speciation in aquatic counterparts **(Rodrigues and Diniz-Filho, 2016; Wiens, 2015a; Wiens, 2015b)**. Higher allopatric speciation leading to higher radiation has been reported in Emydidaen map turtles of south-eastern USA **(Mittermeier et al., 2015)**. It has been observed that rapidly diversifying species are accompanied by higher rates of adaptive evolution as compared to slowly diversifying lineages **(Nevado et al., 2019)**.

Geoemydidae is one of the most diverse Chelonian groups that include the Neotropical wood turtles along with Eurasian pond and river turtles **(Dijk et al., 2012)**. These turtles inhabit freshwater, coastal marine, and tropical forests. Although the closest relative to tortoises, hyper diversity and aquatic lifestyle in Geoemydidae could be associated with higher adaptive selection as detected in genes of CI and CIII of its OXPHOS respiratory system. Emydidae is another megadiverse group of turtles that includes terrapins and marsh turtles. Interestingly, this family also comprises the highly invasive pond slider turtle (*Trachemys scripta*). Human-driven introduction into diverse non-native habitats resulting in novel ecological opportunities for Emydidae species may have led to its higher diversification rate. Associated physiochemical needs could have precipitated in high rates of adaptive selection in CI and CIII of the OXPHOS system of Emydidae. In Platysternidae, adaptive evolution has been exceptionally intriguing as the monotypic family revealed the highest adaptive evolution of all turtles examined. The Big-headed turtle endemic to Southeast Asia and Southern China is a morphological enigma. As carnivores, these turtles possess sharp claws and beak along with a tail, which helps in climbing obstacles in natural habitat **(Pritchard, 1979)**. These energy-expensive morphological and ecological adaptations were congruent with large-scale positive selection detected in its OXPHOS metabolic pathway, across CI, CIII-CV.

Higher adaptive selection in turtles compared to tortoises in Testudinoidea highlights dynamic evolutionary forces acting on closely related species and groups. It underlines the significance of natural pressure pertinent to genomic heterogeneity and warrants a more precise understanding of the nuclear genome and coordinated implication of natural selection. Insights on the evolution of critical functional genes as a response to natural stressors will lead to more efficient ways of conserving these endangered species in the face of the ongoing extinction crisis keeping in mind their natural adaptive potential and its limitations.

## 4. Materials and methods

### 4.1. Dataset analysis

The sequences of the complete mitochondrial genome of 67 species of Testudinoidea were obtained from the NCBI GenBank, comprising: 31 Geoemydidae, 31 Testudinidae, 1 Platysternidae and 4 Emydidae species **(Supplementary Table: ST1)**. The analysis was performed for each PCG fragment, and with concatenated PCGs dataset.

### 4.2. Sequence alignment

The fragments of 12 PCGs sequences encoded on the heavy strand (NAD1, NAD2, COX1, COX2, ATP8, ATP6, COX3, NAD3, NAD4L, NAD4, NAD5, and COB) excluding NAD6 encoded on light strand were extracted from the whole mitogenome and concatenated manually. Multiple sequences alignment was performed with MUSCLE **(Edgar, 2004)** in software MEGA X **(Kumar et al., 2018)**. Concatenated sequences were used for phylogeny estimation and positive selection analyses were implemented independently on each gene.

Nucleotide sequences were translated into AAs, aligned, manually cross-checked with removal of stop codons. Frameshifting indels (insertions or deletions) of 1-2 bp in a few species were manually edited and only fully resolved sites were selected and used for analysis **(Russell and Beckenbach, 2008)**. All sequences were manually edited to preserve the expected reading frame according to the reported indels in turtle mtDNA. Gaps and neighboring ambiguous sites were manually checked.

### 4.3. Codon Saturation test

Codon position-specific patterns of saturation was examined in the concatenated sequences with DAMBE using the test by Xia **(Xia, 2013)**. Estimates of Iss (Index of Substitution Saturation) were used to deduce saturation **(Xia et al., 2003; Xia et al., 2009)**. For codon positions 1 & 2, we obtained estimates indicating minimal substitution saturation for both symmetrical and asymmetrical with significant P-value (<0.001). For the 3^rd^ codon position, we obtained estimates indicating minimal substitution saturation congruent with both symmetrical and asymmetrical tree topology with significant P-value (<0.001). Based on this analysis, the phylogenetic tree retrieved from our dataset that was congruent with the previously reported genetic relationship of turtles and tortoise, indicated negligible saturation in our dataset deeming it fit for further analyses.

### 4.4. Phylogenetic Analyses

The phylogenetic tree was constructed using the best fit model of evolution for each gene separately, and for the concatenated dataset estimated using jModelTest **(Darriba et al., 2012)** with minimal Akaike Information Criterion scores **(Bozdogan, 1987)**. GTR+G+I was identified as the best fit nucleotide substitution model **(Darriba et al., 2012)** and used to generate Maximum Likelihood and Bayesian estimated tree topology.

The Bayesian consensus tree was estimated with the Markov Chain Monte Carlo (MCMC) sampling method using BEAST version 1.7 **(Drummond et al., 2012)**. The MCMC chain was run with four chains for 10^6^ generations, and every 100 samples were taken to estimate the posterior probability distribution. The runs were evaluated in Tracer v. 1.6 (http:/beast.bio.ed.ac.uk/Tracer). The final phylogenetic tree was visualized in FigTree v.1.4.2. (http://tree.bio.ed.ac.uk/software/figtree/). The ML tree was constructed with RAxML using raxmlGUI 2.0, implementing the best-fit model of nucleotide substitution selected as per the Bayesian Information Criterion (BIC). Bootstrap support was obtained by running 1000 pseudoreplicates.

### 4.5. Adaptive selection analyses

The selection and functional divergence analyses were made to fulfil the following objectives: 1) to compare relative patterns and strength of adaptive selection acting on turtles versus tortoise lineages separately, and 2) to identify the patterns and strength of adaptive selection as well as functional divergence across all lineages in Testudinidae

#### 4.5.1. Selective functional divergence in turtles versus tortoises

We performed selection tests to identify sites under positive selection across in turtles and tortoises comparatively, using program TreeSAAP **(McClellan and McCracken, 2001; Woolley et al., 2003)** and CODEML **(Yang, 2007)**.

TreeSAAP measures positive selection on physicochemical AAs properties under the context of phylogeny. The distributions of observed changes inferred from the phylogenetic tree were compared with the expected distribution under a random neutrality model. The turtle and tortoise lineages were analysed separately on TreeSAAP to evaluate lineage specific physicochemical changes indicating selective divergence and comparatively analyse potential implications. We used the default 31 physicochemical properties in our analysis with a sliding window size of 20 codons. To reduce chances of false positives, only those AAs changes were considered, which exhibited radical changes under the categories of magnitude 6 to 8 with significantly positive Z-scores (P < 0.001). AAs substitutions at PSS were inferred from ancestral sequences reconstruction with BASEML in the program TreeSAAP.

To supplement our TreeSAAP analysis, we implement the most widely used program to detect adaptive selection: CODEML in the PAML X software package **(Yang, 2007)**. It detects sites under positive selection under a maximum likelihood framework and codon-based substitution model. It estimates the non-synonymous to synonymous substitution rate ratio (dN/dS, ω) **(Goldman and Yang, 1994; Muse and Gaut., 1994)**. We used Branch site (BrS) analysis to comparatively analyse adaptive selection in turtle as compared to tortoise branches. BrS analysis allows detection of sites under adaptive selection across selective branches/lineages with variable ω in the phylogenetic tree **(Yang et al., 2000; Bielawski and Yang, 2004; Zhang et al., 2005)**. The tree branches are split into a ‘foreground branch’ while the rest of the branches are designated as background branches. The recommended nested model A with null model A (fixed ω=1) was implemented to detect lineage-specific PSS with P < 0.005 **(Yang et al., 2005; Zhang et al., 2005)**. Bayes Empirical Bayes (BEB) **(Yang et al., 2005)** was implemented to calculate PP for site classes to identify PSS in the case of significant LRTs. Separate tests were carried out for each Family in Testudinoidea to compare adaptive evolution in turtles versus tortoises. The branches were labeled based on prior information on phylogeny of Testudinoidea, congruent with fossil evidence **(Crawford et al., 2015)**. Therefore, TreeSAAP and PAML analyses were collectively used to infer functional divergence by analyzing AAs substitutions at PSS identified by CODEML **(McClellan and McCracken, 2001; Woolley et al., 2003)**.

#### Selective functional divergence across Testudinoidea

We again performed selection tests to identify sites under positive selection across Testudinoidea using program TreeSAAP **(McClellan and McCracken, 2001; Woolley et al., 2003)** and CODEML, but executed differently this time. Instead of separate analysis for turtles and tortoises, TreeSAAP was implemented on the entire Testudinoidean tree to estimate AAs substitution-based divergence across the entire Superfamily.

In the program CODEML in the PAML X software package **(Yang, 2007)**, instead of BrS, the Random site analysis was used to detect sites under pervasive adaptive selection, which allows variable ω ratio among sites. Several models were tested in CODEML to detect pattern across all lineages: M0 is the most basic model, which detects the variability of ω among sites rather than explicitly testing for positive selection. Also, we compared nested models with significant Likelihood Ratio Tests (LRTs) to detect PSS comparing the fitness of alternative models versus null models. We finally implemented the null model M8a versus alternative model M8. M8a is characterized by beta (β) with fixed ω while M8 detects PSS characterized with β and variable ω **(Yang et al., 2000; Swanson et al., 2003)**. BEB **(Yang et al., 2005)** was implemented to calculate PP for site classes to identify PSS in the case of significant LRTs.

#### Structural analyses with Protein modeling

To understand the structural and functional significance of adaptive selection in mitochondrial protein complexes of the OXPHOS system, PSS were mapped onto the protein sequences of each subunit of complex I, III, IV, and V were aligned with homologous protein subunits available in NCBI GenBank database. Protein homology models of each subunit were then predicted from the best hit template in the SWISS-MODEL server **(Supplementary Table: ST7)**. An overall structure of each OXPHOS complex was created by aligning individual subunits with the corresponding complex’s template. We detected the residue location of PSS on proteins and their associated physicochemical characteristics with TMHMH v2.0 and solved crystallized structures. Graphical depiction of PSS across the OXPHOS system was visualized using PyMOL software **(Schrodinger, 2017)**.

## 5. Conclusion

Our study furnishes compelling evidence for different selective pressure acting on closely related species through their adaptive mitochondrial genome. We found stronger positive selection in the turtle mitochondrial genome compared to limited positive selection in tortoises. The study highlights the vital role of contrasting environments in shaping the adaptive evolutionary changes in the OXPHOS respiratory system in turtles and tortoises. This may be closely associated with greater habitat diversification, higher energy demanding physiochemistry, morphological adaptations and a higher rate of speciation in turtles as compared to tortoises. Our study revealed a complex pattern of mitochondrial DNA evolution across Testudinoidea, indicating predominant adaptive destabilizing convergent selection in turtles compared to stabilizing divergent selection in tortoises. Moreover, in turtles, we found numerous unique sites across gene complexes under positive selection in Platysternidae. This indicated that the mitochondrial OXPHOS genes were actively involved in determining functional morphological and physiological adaptations among lineages to suit their different environmental challenges.

The study illustrated several adaptively selected sites detected in structurally and functionally significant regions of the OXPHOS system. This paves way for further research into biochemical implications of genetic substitution in closely related species. Our study provides novel insights into contrasting adaptive potentials of one of the most archaic and evolutionarily enigmatic yet threatened reptile groups, which may have enabled historical successful eco-physiological diversification of these Testudinoidean species into their diverse environmental habitats.

## Acknowledgments

S.S. was provided fellowship by the University Grant Commission, Ministry of Human Resources and Development, Government of India. We acknowledge the support provided by the Director and Dean, Wildlife Institute of India, and the entire team of Wildlife Forensic and Conservation Genetics Cell.

## Competing interests

The author(s) declare no competing interests.

## Data accessibility

All supplementary data cited in the main text and additional information has been uploaded online as supporting information.

## Author contributions

The study was conceived and designed by A.K. and S.K.G. Computational analysis, data interpretation and manuscript writing were carried out by S.S. and A.K. Finalization and refining of the manuscript were done by J.R. and S.K.G. All authors have read and approved the final version of the paper.

## Notes

### Competing Interest Statement

The authors have declared no competing interest.

